# Optimizing CRISPR/Cas9 mutagenesis in Drosophila da neurons to avoid cytotoxicity

**DOI:** 10.1101/2025.10.03.680356

**Authors:** Vinicius N. Duarte, Viana Najafi, Katherine L. Thompson-Peer

## Abstract

Genetic perturbations are one of the great strengths of the model organism *Drosophila melanogaster*, with approaches such as classical mutagenesis and RNA interference enabling a wealth of biological discoveries. A more recent approach for altering gene expression is CRISPR/Cas9-based mutagenesis, but as with any new tool, its use must be optimized. High expression of Cas9 has been shown to cause cytotoxicity in some cell types. Here, we show that Cas9 expression alone causes cytotoxicity in the dendritic arborization (da) neurons that are widely used to study neuronal development and regeneration. We then systematically evaluate alternative Cas9 transgenes designed to lower total Cas9 expression, called uCas9 transgenes. We show that expression of these uCas9 transgenes results in little to no cytotoxicity to da neurons. Lastly, we demonstrate the ability of uCas9 transgenes to effectively and specifically gene edit in da neurons. Thus, we expand the toolkit of genetic perturbations available to researchers working with Drosophila da neurons or other cell types suceptible to cytotoxicity due to high expression of Cas9.

## Introduction

The CRISPR (clustered regularly interspaced short palindromic repeats) gene editing platform has revolutionized the research community’s ability to probe gene function^1^. As with any new scientific tool, its use in model organisms and cultured cells has required optimization. Ectopic expression of the Cas9 protein, an endonuclease that induces DNA damage with specificity from guide RNA molecules, can cause cellular toxicity, as reported by multiple groups ^2–4^. Interestingly, this is not dependent on endonuclease activity as expression of a nuclease-dead Cas9 protein, widely used to increase gene expression via its fusion to transcriptional activators, also results in cellular toxicity^2^.

The dendritic arborization (da) neurons of *Drosophila melanogaster* are a powerful system to examine neuronal development and regeneration^5^. There are four different classes of da neurons (type I-IV) based on increasing dendritic complexity. Class IV da neurons have highly branched dendrite arbors, and they continue growing throughout larval development. Due to the large and dynamic nature of the class IV dendrite arbor, class IV neurons are often the most sensitive to manipulations that affect dendrite development and outgrowth. High expression of Cas9 protein via the Gal4/UAS system in the class IV da neuron subtype results in severe cytotoxicity as evidenced by reductions in dendrite density^6^. Because of this cytotoxicity, it has been challenging to use CRISPR/Cas9 manipulations to study gene function during dendrite development and regeneration in da neurons.

Some groups have taken the approach to prevent this dendrotoxicity by creating enhancer-driven Cas9 expression vectors that reduce levels of Cas9 protein^6^. However, this approach has some limitations. First, genetic perturbations in different tissue or cell types would require new cloning and transgenesis procedures that could be costly, time-consuming, and technically demanding. Second, enhancer driven Cas9 is likely expressed earlier than Gal4/UAS-mediated Cas9^7^. This may not be favorable for studies seeking to limit developmental confounds associated with gene knockout too early in development. Third, enhancer-driven Cas9 expression levels may differ across enhancers, creating uncertainty that could require further fine-tuning of experimental design. That is, after generating new enhancer-driven Cas9 expression vectors and creating new fly stocks with transgenesis, cytotoxicity may still be present depending on the strength of the enhancer.

An alternative approach to titrating Cas9 expression levels may be achieved with UAS-Cas9 transgenes containing upstream open reading frames (uORFs) of varying lengths that are inversely correlated with Cas9 expression^8,9^. These transgenes are referred to as uCas9 and initially Port et al (2020) generated a series of six lines with different uORF lengths ranging from 33 bp to 714 bp. In Drosophila wing imaginal discs, Cas9 immunostaining revealed a gradual decrease in Cas9 expression with increasing uORF length. Correspondingly, Cas9-induced apoptotic cell number gradually decreased, and target gene expression gradually increased. Thus, this approach successfully tunes Cas9 expression levels, prevents Cas9-mediated cytotoxicity and maintains Cas9 gene editing efficiency in the wing imaginal disc^8^. We hypothesized that these uCas9 transgenes may present a suitable alternative to CRISPR-based mutagenesis in da neurons, particularly for studies seeking to circumvent the limitations associated with the enhancer-driven Cas9 approach.

Here, we first validated Cas9-induced dendrotoxicity in class IV da neurons using a transgene with no uORF, which resulted in high Cas9 expression and dendrotoxicity. We also assessed Cas9-induced dendrotoxicity in two other widely studied da neuron types: class I and class III da neurons. We then determined which of three different uCas9 transgenes with varying uORFs were most suitable for mitigating cytotoxicity in each neuron type. These were uCas9 (M), uCas9 (L), and uCas9 (VL) corresponding to medium, low, and very low expression of Cas9. Lastly, we confirmed CRISPR-based gene editing efficiency when targeting GFP in class I da neurons. Thus, we showcase an alternative approach to minimizing Cas9-associated cytotoxicity via the use of uCas9 transgenes and establish the most optimal uCas9 transgene for CRISPR-based mutagenesis in Drosophila da neurons.

## Results

To assess Cas9-induced dendrotoxicity in Drosophila da neurons, we imaged late third instar larvae expressing UAS-driven Cas9 in class IV, class I, or class III da neurons using the cell-type specific drivers ppk-Gal4, 2.21-Gal4, and 19.12-Gal4, respectively (Figure 1A). We call the expression of Cas9 without any uORF as “Cas9(H)” for high expression. Expression of this Cas9 transgene caused noticeable changes to dendrite morphology in class IV ddaC neurons (**Figure 1A**, top row, orange tracing relative to black). Quantifications of branch number and total dendrite length confirm that both measures are significantly reduced in Cas9 (H)-expressing class IV ddaC neurons (**Figure 1B**, top row). We extended this assessment to class I ddaE neurons, which also show a significant reduction in branch number for those expressing Cas9 (H) (**Figure 1B**, middle row). Lastly, expression of this Cas9 transgene in class III ddaA neurons did not affect either branch number or total dendrite length (**Figure 1B**, bottom row). The lack of dendrotoxicity in class III da neurons is likely due to the late expression of the Cas9 in this cell type, since the 19.12-Gal4 driver is not active until the late 2^nd^ instar stage of Drosophila development. This provides further support to the model that Cas9 at too high a level results in dendrotoxicity.

**Figure 1:**
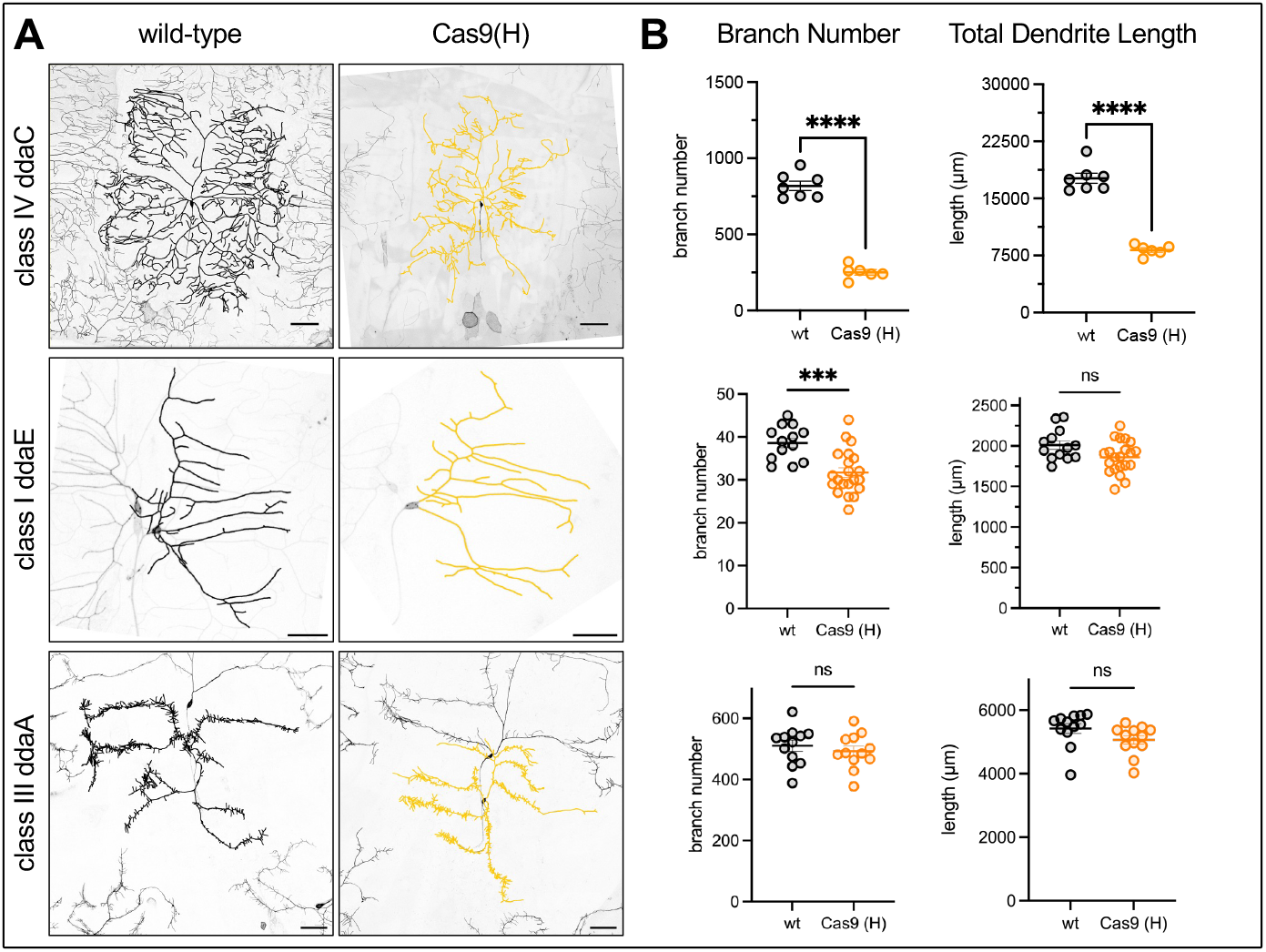
Cas9 expression in da neurons causes dendrotoxicity. (A) presentative images of class IV ddaC (top row), class I ddaE (bottom row), or ass III ddaA in third instar larvae of wild-type (left column, black overlay) or Cas9(H)-expressing (right column, orange overlay) neurons. Traces are overlaid onto neurons of interest for ease of visualization and all images are rotated axon side down. Scale bars = 100µm for class IV ddaC images and 50µm for class I ddaE and class III ddaA images. (B) Quantifications of branch number and total dendrite length with statistical comparisons between wild-type and Cas9(H)-pressing neurons. Comparisons were made using Welch’s t-test. Average and error are mean ± SEM. ^*^ p < 0.05, ^**^ p – 0.01, ^***^ p < 0.001.

To titrate levels of Cas9 protein, we expressed uCas9 transgenes with varying uORF lengths in class IV ddaC, class I ddaE, and class III ddaA neurons (**Figure 2A**). In class IV ddaC neurons, quantifications of branch number are significantly disrupted in Cas9 (H)-expressing neurons. Branch numbers largely return to normal in uCas9 (M), uCas9 (L), and uCas9 (VL)-expressing neurons relative to wild type, with uCas9M and uCas9VL surprisingly having more branches than wild-type (**Figure 2B**, top row). While Cas9 (H) has a defect in total branch length compared to wild-type, total dendrite length was normal for all of the uCas9 transgenes (data not shown). In class I ddaE neurons, quantifications of branch number are again significantly reduced in Cas9 (H)-expressing neurons, but normal in all of the uCas9 transgenes (**Figure 2B**, middle row). Total dendrite length was significantly reduced in uCas9 (M)-expressing ddaE neurons (data not shown). In class III ddaA neurons, no changes in branch number (**Figure 2B**, bottom row) or total dendrite length (data not shown) were observed with any of the transgenes tested, similar to the absence of dendrotoxicity observed in Fig 1. Altogether, these suggest that uCas9 (L) is the optimal transgene to express in class IV and class I da neurons, while Cas9 (H) is suitable for expression in class III da neurons.

**Figure 2:**
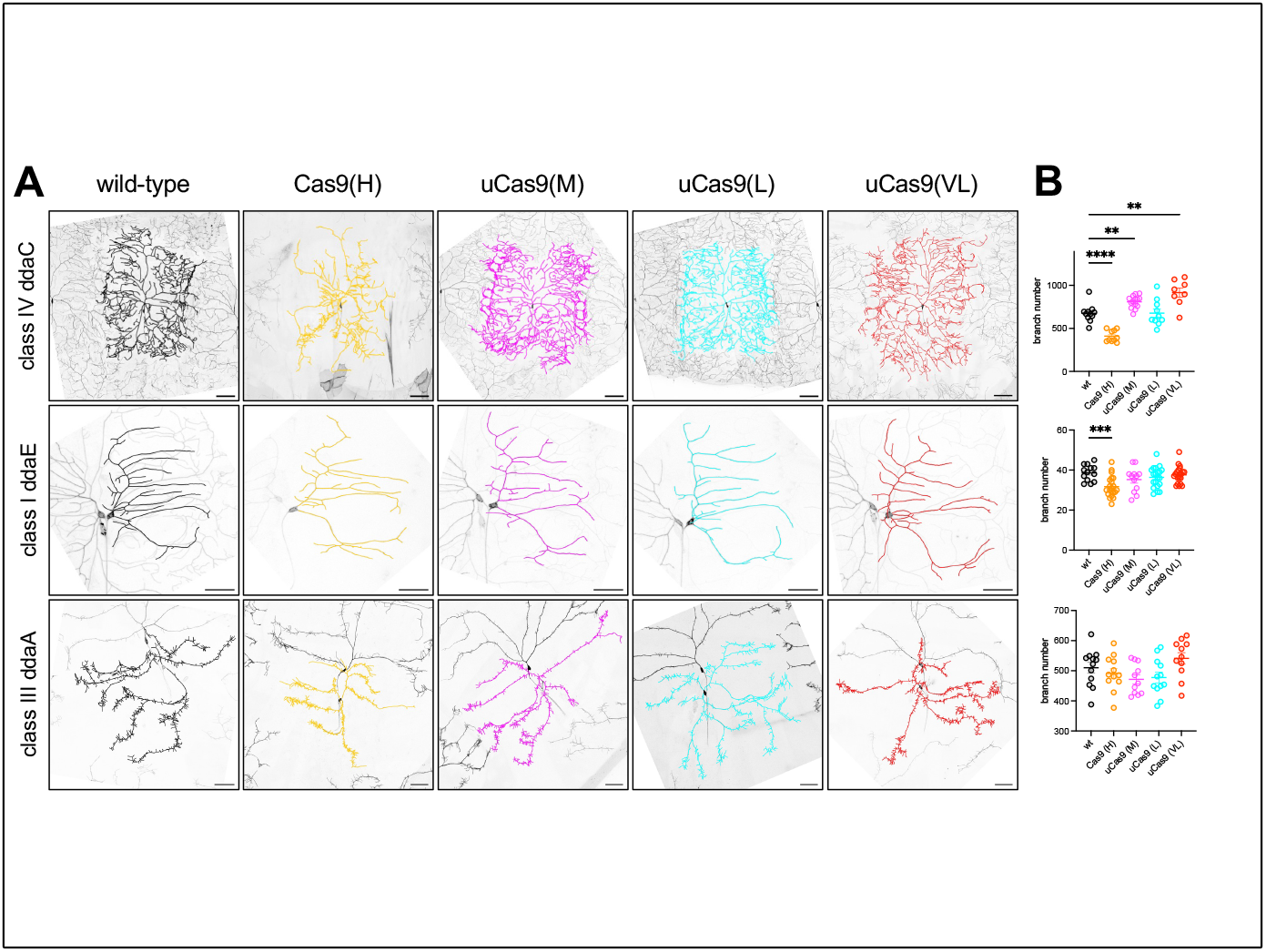
uCas9 transgenes reduce dendrotoxicity. (A) Representative images class IV ddaC (top row), class I ddaE (middle row, or class III ddaA (bottom w) neurons in third instar larvae as wild-type (first column, black overlay), Cas9 (H)-expressing (second column, orange overlay), uCas9 (M)-expressing (third column, magenta overlay), uCas9 (L)-expressing (fourth column, cyan overlay), uCas9 (VL)-expressing (fifth column, purple overlay). Scale bars = 100µm for ass IV ddaC images and 50µm for class I ddaE and class III ddaA images. (B) uantifications of branch number and total dendrite length with significant statistical comparisons shown. Comparisons were made using Brown-Forsythe and Welch’s ANOVA with Dunnett multiple comparison correction. Average and error are mean ± SEM. ^*^ p < 0.05, ^**^ p – 0.01, ^***^ p < 0.001.

Lastly, we wished to assess gene editing efficiency using the identified uCas9 (L) transgene. To do this, we drove uCas9 (L) and EB1::GFP in class I da neurons and ubiquitously expressed (1) a single unique gRNA for QUAS (control), (2) a single unique gRNA for EGFP, or (3) triple gRNAs for GFP, EGFP, and BFP with Ubi-BFP::nls as a co-CRISPR reporter. In animals with control gRNA for QUAS, a clear EB1::GFP signal was detected, indicating that no gene editing had occured (**Figure 3A**). In animals with gRNA targeting EGFP, no EB1::GFP signal was detected (**Figure 3A**). Denticle belts and larval body edges were used as guideposts to find the expected class I da neuron location. In animals with triple gRNAs targeting GFP, EGFP, and BFP, along with Ubi-BFP::nls as an internal control, no EB1::GFP signal was detected (**Figure 3A**). Importantly, Ubi-BFP was still expressed in surrounding cells with no uCas9 (L). All segments imaged with the control gRNA for QUAS had a clear EB1::GFP signal, while no segments imaged for the other two groups did (**Figure 3B**). These data provide strong evidence for the efficacy and specificity of uCas9 (L) mutagenesis in Drosophila da neurons.

**Figure 3:**
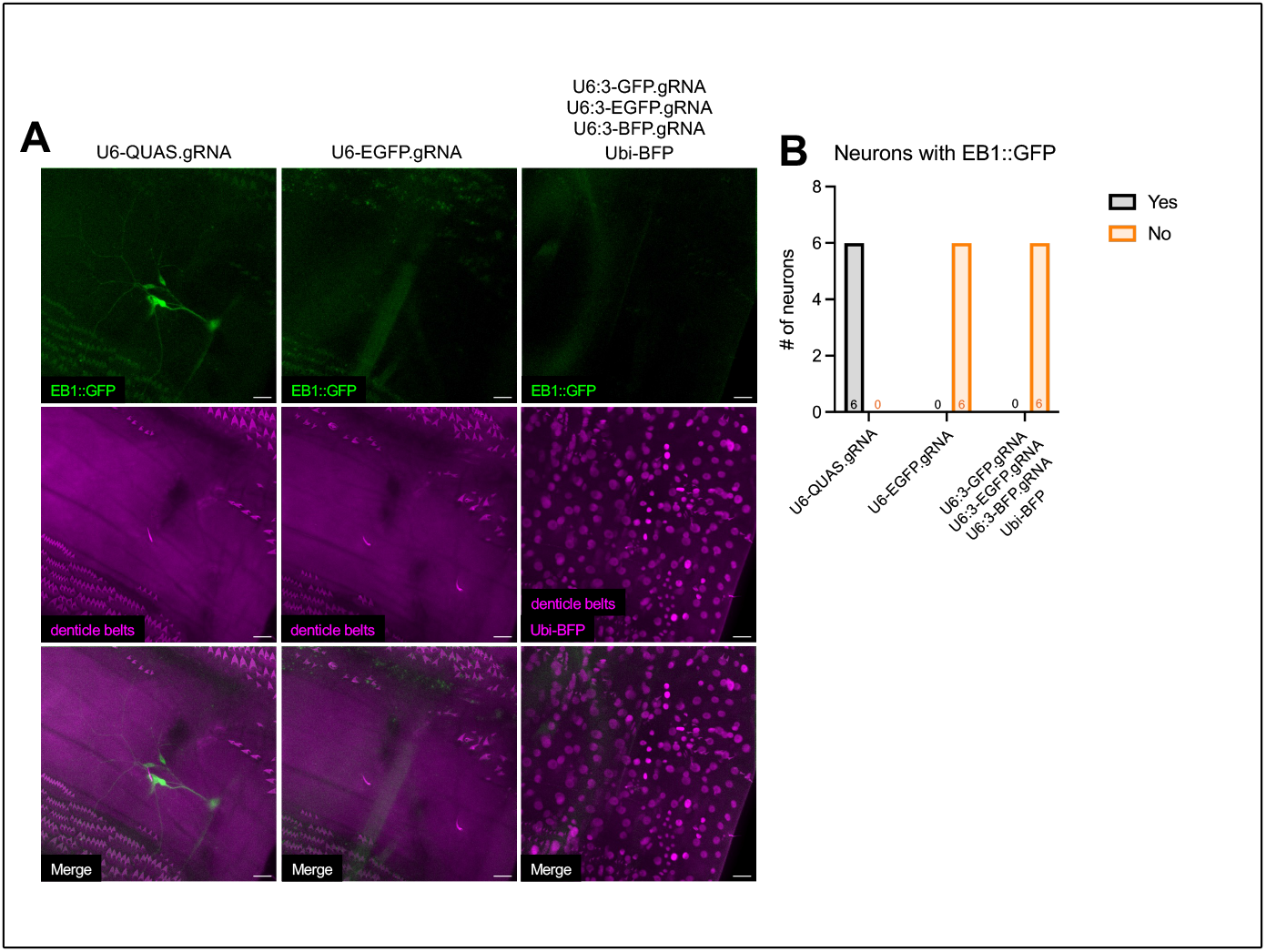
uCas9 transgenes are capable of gene editing in Drosophila da nrurons. (A) Representative images of class I da neurons with pdm-Gal4 driven UAS-EB1::GFP and UAS-uCas9(L). gRNAs were ubiquitously driven to target QUAS (first column, control), a single EGFP.gRNA (second column), or triple NAs targeting GFP, EGFP, and BFP (third column, with Ubi-BFP). Scale bars = µm. (B) Qualitative assessment of EB1::GFP expression for all genotypes. Three neurons were imaged per animal, two animals per genotype. Fisher’s exact test yielded a p = 0.0002.

## Discussion

This study offers an alternative approach to CRISPR/Cas9-mediated gene editing using the Gal4/UAS system and uCas9 transgenes that mitigate cytotoxicity. In addition to class IV da neurons, we extend findings of cytotoxicity to the more stable class I da neurons. We find no evidence of cytotoxicity in class III da neurons, perhaps due to the late expression of the Gal4 driver used to drive in that cell type. After systematic analysis of various uCas9 transgenes, we identified uCas9 (L) as the most optimal transgene for da neurons. Finally, we show that uCas9 (L) mutagenesis is effective and specific. Since the Gal4/UAS system is widely used by Drosophila researchers for spatiotemporal transgene expression, UAS-uCas9 transgenes should prove useful for those seeking to incorporate CRISPR/Cas9 mutagenesis in their studies.

## Materials and Methods

### Fly Stocks and Drosophila Culture

Class I da neurons were labeled by UAS-CD4::tdTomato under 2.21-Gal4^10^, class III da neurons were labeled by UAS-CD4::tdGFP under 19.12-Gal4 with glial repression under repo-Gal80^11,12^, and class IV da neurons were labeled by ppk-CD4::tdGFP^7^. Pdm-Gal4 was used in Figure 3 (gift from Chun Han). Cas9 transgenes used in this study include UAS-Cas9 (H) (Bloomington #54595), UAS-uCas9 (M) (Vienna #340002), UAS-uCas9 (L) (Vienna #340003), and UAS-uCas9 (VL) (Vienna #340004)^8^. To drive Cas9 expression in da neurons, 2.21-Gal4/pdm-Gal4 was used for class I da neurons, 19.12-Gal4 was used for class III da neurons, and ppk-Gal4^13^ was used for class IV da neurons. We used the following Bloomington Drosophila Stock Center gRNA lines to determine uCas9 (L) gene editing efficiency: 67539 (QUAS.gRNA), 79393 (EGFP.gRNA) and 92748 (GFP.gRNA, EGFP.gRNA, BFP.gRNA, Ubi-BFP::nls).

Adult fly crosses were used for 2-4h synchronized egg lays and larvae were reared at 22.5°C in humidified incubators until the third instar stage. Identical culture conditions were used within datasets.

### Live Imaging of Drosophila Larvae

Third instar larvae were mounted one by one on an agarose pad atop a glass slide. Dollops of vacuum grease were placed around the agarose pad, and a drop of glycerol was placed onto the larva. Coverslips were pressed onto the vacuum grease, with the larva oriented dorsal side up and tape was used to secure the coverslip. Animals were imaged using a Zeiss LSM980, Zeiss LSM900, or Zeiss LSM700 microscope with a 20x objective. Z-stacks containing the full dendrite arbor were acquired. For class IV dendrite arbors, the tiles function was used to ensure acquisition of the entire arbor.

### Quantification and Statistical Analyses

Image processing and analysis was done in Fiji/ImageJ. Z-stacks were maximum intensity projected and median filtered for clarity. Dendrites arbors were traced using the Simple Neurite Tracer plugin^14,15^. Branch number and total dendrite length were acquired from Simple Neurite Tracer. GraphPad Prism was used to generate graphs and perform statistical analyses. A significance threshold of p < 0.05 was used for all comparisons, and detailed information on specific statistical tests used can be found in figure captions.

## Acknowledgements

This study was supported by R00NS097627 and startup funds from the University of California, Irvine (to KTP). VND received support from T32MH119049 and the UCI School of Biological Sciences ODEI Postdoctoral Excellence Fellowship. This study was made possible in part through access to the Optical Biology Core Facility of the Developmental Biology Center, a shared resource supported by the Cancer Center Support Grant (CA-62203) and NIH-S10OD032327-01. Transgenic fly stocks obtained from the Bloomington Drosophila Stock Center (NIH P40OD018537), the Vienna Drosophila Resource Center (VDRC, www.vdrc.at), and Dr. Chun Han were used in this study. KTP is a fellow of the Hellman Family Foundation and the Rose Hills Foundation.

